# PCSK9 and High-Fat Diet Synergistically Induce Neurovascular Dysfunction and Neuroinflammation

**DOI:** 10.1101/2025.10.29.685380

**Authors:** Mengqi Zhang, Anudari Letian, Gregory A Mohl, Brynn Kroke, Kajsa Arkelius, Claire D Clelland, Emily L Goldberg, Neel S. Singhal

**Affiliations:** Department of Neurology, University of California- San Francisco, San Francisco, CA, USA; Department of Physiology, University of California- San Francisco, San Francisco, CA, USA; Department of Neurology, Memory and Aging Center, University of California- San Francisco, San Francisco, CA, USA; Chan-Zuckerberg Biohub, San Francisco, CA, USA; San Francisco Veterans Affairs Medical Center, University of California- San Francisco, San Francisco, CA, USA

**Keywords:** Neuroinflammation, PCSK9, Hypercholesterolemia, Lipid droplet–accumulating microglia, Cerebral small vessel disease

## Abstract

Cerebral small vessel disease (CSVD) is strongly linked to metabolic risk factors and represents a major cause of vascular cognitive impairment and dementia. The interactions of genetic and environmental risk factors driving cerebrovascular pathology in metabolic syndrome are poorly understood. Here, we characterize neuroinflammatory and neurodegenerative phenotypes in a mouse model of metabolic syndrome with atherosclerosis induced by hepatic proprotein convertase subtilisin/kexin type 9 (PCSK9) overexpression combined with high-fat diet (HFD). PCSK9+HFD mice exhibit hallmark features of CSVD including vascular rarefaction, impaired neurovascular coupling, blood-brain barrier disruption, white matter injury, neuronal loss, and cognitive deficits. Notably, we identify lipid-droplet accumulating microglia (LDAM) as a distinct cellular phenotype that emerges in response to metabolic stress and correlates with cerebrovascular dysfunction. Three-dimensional light sheet microscopy reveals widespread vascular network disruption. Immunophenotyping demonstrates that microglia in PCSK9+HFD group exhibit enhanced phagocytic activation and ramification complexity yet accumulate perivascular amyloid-β, suggesting impaired clearance capacity. Importantly, we observed vascular amyloid-β deposition in wild-type mice without genetic Alzheimer’s disease mutations, suggesting that metabolic stress contributes to cerebrovascular amyloid pathology. PCSK9+HFD mice displayed recognition memory deficits and increased anxiety-like behavior. Our findings establish that severe hypercholesterolemia accelerates CSVD pathogenesis, and identify LDAM as a distinct pathological feature linking systemic metabolic syndrome to cerebrovascular dysfunction and cognitive impairment.

## Introduction

Cerebral small vessel disease (CSVD) affects over 50% of the population over 65 years worldwide and is strongly associated with metabolic syndrome, which includes hypertension, hyperlipidemia, insulin resistance, and central obesity[1, 2]. CSVD pathology is strongly associated with lacunar stroke, vascular cognitive impairment and dementia (VCID)[3]. CSVD pathology includes arteriosclerosis, vascular rarefaction, blood-brain barrier (BBB) disruption, gliosis, and white matter injury, and ultimately leads to neuronal dysfunction and atrophy[1, 4]. Recent human studies have also found deficits in neurovascular coupling and cognition in patients with MRI hallmarks of CSVD who are cognitively unimpaired by clinical criteria[5]. Despite its prevalence and significant impact on stroke risk and cognitive function, therapeutic options remain limited, largely due to an incomplete understanding of the molecular mechanisms directly linking metabolic risk factors to cerebrovascular pathology[3].

Metabolic syndrome, characterized by obesity, hyperlipidemia, insulin resistance and hypertension dramatically increases CSVD risk[6, 7]. Brain neuroinflammatory changes such as microglial activation, white matter injury and loss of blood-brain barrier integrity are strongly associated with metabolic syndrome[8]. Hypercholesterolemia, a key component of metabolic syndrome, is strongly associated with neuroinflammatory markers, cognitive impairment and dementia in humans[9]. Animal models of hypercholesterolemia demonstrate neuroinflammation precedes neurodegeneration[10], [11], [12], [13]. Mice globally lacking a key cholesterol receptor, low-density lipoprotein receptor (*LdlR)*, which develop marked elevations in circulating cholesterol due to impaired LDL uptake, have increased glial fibrillary acidic protein (GFAP) immunoreactivity and microglial density in the hippocampus and deficits in hippocampal-dependent learning tasks [14] [15].

However, current CSVD animal models present important limitations. Most models utilize single-factor approaches. Previous work in animal models of CSVD presents mild phenotypes with short-term hypercholesterolemia or relies on genetic alterations that do not capture the gene-environment interactions characteristic of human metabolic syndrome-related CSVD [16]. In addition, models utilizing carotid occlusion or monogenic forms of CSVD may involve different pathophysiological mechanisms than those common in human CSVD related to vascular risk factors. To address this gap, we developed a model combining genetic risk with environmental factors to better recapitulate the multifactorial etiology and synergistic metabolic effects underlying human metabolic syndrome-associated cerebrovascular disease.

Proprotein convertase subtilisin/kexin type 9 (PCSK9) plays a crucial role in cholesterol metabolism by promoting the degradation of LDLRs, thereby reducing LDL cholesterol clearance from the circulation[17]. Gain-of-function mutations in *PCSK9* cause familial hypercholesterolemia in humans[18], and PCSK9 inhibitors have emerged as effective LDL-lowering therapeutics[19]. Beyond its role in cholesterol metabolism, PCSK9 exerts pro-inflammatory effects on vascular cells and may directly influence neural tissue[20-22]. While hepatic PCSK9 overexpression in mice via adeno-associated virus (AAV) delivery has been used to model atherosclerosis in the periphery[23], its effects on the cerebrovascular system remain unexplored. This approach enables sustained, severe hypercholesterolemia in adult mice on a wild-type genetic background. When combined with HFD, this model exhibits marked metabolic dysregulation consistent with features of human metabolic syndrome, including obesity. Understanding these gene-environment interactions is essential to elucidating mechanisms underlying metabolic syndrome-related cerebrovascular disease.

In this study, we systematically evaluated the pathological, neurovascular, and behavioral consequences of liver-specific PCSK9 overexpression combined with HFD in C57BL/6J mice as a novel model of metabolic syndrome-related CSVD. We compared the interactions between PCSK9 overexpression and diet to distinguish their individual contributions to cerebrovascular pathology. Our findings demonstrate a novel pathological mechanism whereby PCSK9 overexpression with HFD synergistically induces microglial lipid accumulation and promotes a comprehensive spectrum of CSVD-like pathology, including neurovascular dysfunction, BBB disruption, neuroinflammation, and cognitive impairment. Our findings establish a mechanistic link between PCSK9-mediated metabolic dysfunction and cerebrovascular disease, providing a valuable platform for therapeutic development in age-related cognitive decline and VCID.

## Methods

### Animals and Experimental Design

Male C57BL/6J mice (7 weeks old) were obtained from Jackson Laboratory and housed under standard specific pathogen-free conditions (22±2°C, 12-hour light/dark cycle). Following one-week acclimatization, cohorts of mice were randomly assigned to four groups (n=10 per group: Control, PCSK9, HFD and PCSK9+HFD). AAV8 vectors encoding mouse PCSK9-D377Y gain-of-function mutant (2×10^11^ genome copies in 100 μL PBS, Addgene) were administered to PCSK9 and PCSK9+HFD groups via retro-orbital injection under brief isoflurane anesthesia (2% induction, 1.5% maintenance). Control and HFD groups received equivalent volumes of vehicle. Dietary interventions were initiated immediately post-injection: HFD and PCSK9+HFD groups received high-fat diet (Clinton/Cybulsky Diet With 60 kcal% Fat and 1.25% Added Cholesterol, Research Diets), while Control and PCSK9 groups were maintained on standard rodent chow (Teklad Global 16% Protein Rodent Diet; irradiated; no added cholesterol; ≥16% crude protein, ≥3.5% crude fat, ≤5.0% crude fiber). All diets were continued for 20 weeks. All procedures were approved by the Institutional Animal Care and Use Committee at UCSF and San Francisco Veterans Affairs Medical Center.

### Serum collection and processing

At the study endpoint (week 20), mice were anesthetized with isoflurane, and blood was collected by cardiac puncture into additive-free serum tubes. Samples were allowed to clot for 30 min at room temperature and centrifuged at 3,000 × g for 15 min at 4 °C. Serum was aliquoted and stored at −80 °C until assay. Total cholesterol was measured using an enzymatic colorimetric assay on a microplate reader according to the manufacturer’s instructions.

### Immunofluorescence Staining

At experimental endpoint, mice were deeply anesthetized with isoflurane. For visualization of perfused vasculature, DyLight 649-conjugated Lycopersicon esculentum lectin (Vector Laboratories, 10 mg/kg body weight) was administered via cardiac perfusion 5 minutes before euthanasia to label functional vessels. Subsequently, mice were transcardially perfused with 0.9% NaCl followed by 4% paraformaldehyde in PBS. Brains were extracted, post-fixed in 4% PFA overnight at 4°C, and cryoprotected in 30% sucrose in PB for 48 hours until sinking. Using a cryostat, brains were sectioned coronally. Sections were stored at -20°C in cryoprotectant solution until processing.

For standard immunofluorescence, free-floating sections were washed with PBS (3×10 minutes) to remove cryoprotectant, followed by permeabilization with 1% Triton X-100 in PBS for 20 minutes, then washed twice with 0.3% PBST (5 minutes each). Sections were blocked with 5% normal donkey serum in 0.3% PBST for 1 hour at room temperature. Primary antibodies diluted in 0.3% PBST were applied overnight at 4°C: rabbit anti-Iba1 (1:500, Wako), chicken anti-NeuN(1:1000, Millipore), chicken anti-GFAP (1:500, Abcam), rabbit anti-CD68 (1:500, Abcam), mouse anti-ZO-1 (1:200, Invitrogen), and mouse anti-amyloid-β (1:200, Santa Cruz). For co-staining of Iba1 with CD68, goat anti-Iba1 (1:500, Abcam) was used instead of rabbit anti-Iba1 to avoid species cross-reactivity. Prior to applying mouse primary antibodies, sections were treated with a mouse-on-mouse blocking kit (Vector Laboratories) according to the manufacturer’s instructions to minimize non-specific binding.

For lipid droplet accumulation analysis, separate protocols were employed. For Iba1/perilipin-2 co-staining, sections were blocked with 5% normal donkey serum in PBS (without Triton X-100) for 1 hour, then incubated overnight at 4°C with goat anti-Iba1 (1:500, Abcam) and rabbit anti-perilipin-2 (1:200, Abcam). After primary antibody incubation and PBS washes, appropriate Alexa Fluor-conjugated secondary antibodies (1:500) were applied for 1 hour at room temperature.

After all staining procedures, sections were washed with PBS (3×10 minutes), mounted on glass slides and coverslipped with ProLong Gold antifade mounting medium (Vector Laboratories).

Images were acquired using either a Zeiss AXIO Observer.A1 inverted fluorescence microscope or a CSU-W1 SORA spinning disk confocal microscope, depending on the specific analysis requirements. For quantification, 3 non-consecutive sections per animal were analyzed. Three randomly selected fields per section were evaluated for each brain region (cortex, corpus callosum, external capsule, striatum, hippocampus). Vascular parameters, immunoreactive areas, and cell counts were quantified using ImageJ software. Three-dimensional reconstructions were generated using Imaris software (Oxford Instruments) for spatial relationship analysis.

### Luxol Fast Blue Staining

White matter integrity was assessed using Luxol Fast Blue (LFB) staining. Mounted brain sections were incubated in 0.1% LFB solution (StatLab) at 60°C for 2-4 hours until optimal staining was achieved. After cooling to room temperature, sections were differentiated using 0.05% lithium carbonate solution and 70% ethanol until white matter appeared distinctly blue against pale gray matter. Sections were then dehydrated through ascending ethanol concentrations (70%, 95%, 100%), cleared in xylene, and coverslipped. Images were acquired using a Zeiss AXIO Observer.A1 microscope. Optical density was measured in corpus callosum, external capsule, and striatum from three non-consecutive sections per animal (3 fields per section) using ImageJ software. Values were normalized to background and expressed as relative optical density.

### Laser Speckle Contrast Imaging

Neurovascular coupling was assessed using laser speckle contrast imaging (LSCI). Mice were anesthetized with isoflurane (1.5% maintenance) and placed in prone position with body temperature maintained at 37°C via heating pad. Following midline scalp incision to expose the skull, the intact skull surface was kept moist with saline. A laser speckle contrast imager (RFLSI III, RWD Life Science) with 785 nm laser illumination was positioned 10 cm above the skull. After stabilization, baseline cerebral blood flow (CBF) was recorded before initiating contralateral whisker stimulation (5 Hz, 30 seconds). Regions of interest (ROIs) were defined over the barrel cortex corresponding to stimulated whiskers (as indicated by black ovals in images). Three trials were performed per animal with 5-minute inter-trial intervals. CBF responses were quantified as percentage change from baseline: [(Mean perfusion during stimulation - Mean perfusion at baseline) / Mean perfusion at baseline] × 100%. Values were averaged across three trials per animal. For time-course analysis, CBF changes were calculated at 2-second intervals throughout the recording period.

### Light Sheet Microscopy

Whole-brain clearing and imaging were performed by LifeCanvas Technologies. Brains previously perfused with DyLight 649-lectin were processed using the SHIELD tissue preservation and SmartClear Pro clearing system according to manufacturer’s protocols. Briefly, fixed brains underwent delipidation and refractive index matching to achieve optical transparency. Cleared samples were immunolabeled with anti-Iba1 antibody using SmartLabel. Three-dimensional imaging was performed using SmartSPIM light sheet microscopy with 4× detection objective at 1.8 × 1.8 × 4 μm voxel resolution. Image stacks were processed and 3D reconstructions generated using Imaris 10.2 (Oxford Instruments) for visualization of vascular-microglial spatial relationships.

### Behavioral Testing

#### Open field test

Spontaneous exploration and anxiety-related behavior were assessed in an open field arena (40 × 40 × 35 cm). Animals were habituated to the testing room for ≥30 min before testing. The arena was uniformly illuminated and recorded from above with an overhead camera. A center zone (16 × 16 cm) was predefined in ANY-maze. At the start of each trial, a mouse was gently placed in the center and allowed to explore freely for 5 min. Locomotor trajectories were tracked automatically. The primary outcomes were time spent in the center zone and time spent in the corner zones (thigmotaxis). ANY-maze software tracked movement patterns and quantified total distance traveled, center zone time, corner zone time, and average velocity. The arena was cleaned with 70% ethanol and dried between trials to remove olfactory cues.

### Novel object recognition (NOR)

To assess recognition memory, the NOR test was conducted following established protocols. In the habituation phase, mice freely explored the empty arena for 5 minutes. During the familiarization phase, mice were exposed to two identical objects and allowed to explore for 10 minutes. Animals were introduced facing the arena wall to ensure unbiased initial exploration. After a 1-hour inter-trial interval in the home cage, the test phase commenced with one familiar and one novel object for 10 minutes of exploration. The arena and objects were cleaned with 70% ethanol between trials. Object investigation was defined as nose orientation toward the object within 2 cm distance, including sniffing or touching. ANY-maze provided exploration times for the familiar (T_familiar) and novel (T_novel) objects, and the discrimination index was calculated as (T_novel − T_familiar) / (T_novel + T_familiar).

### Statistical Analysis

Data were analyzed using GraphPad Prism version 10.2.3. Normality was assessed using the Shapiro–Wilk test, and homogeneity of variance with the Brown–Forsythe test. For datasets that met normality and homoscedasticity assumptions, one-way ANOVA was used for single-factor (≥3 groups) comparisons, followed by Tukey’s multiple comparisons test. When two independent factors were present, a two-way ANOVA was performed with Šidák’s post-hoc tests. Welch’s ANOVA with Dunnett’s T3 post-hoc was applied when variances were heterogeneous. Data are presented as mean ± SEM. Statistical significance was set at P<0.05.

## Results

### Hypercholesterolemia induced by PCSK9 overexpression and HFD drives neurovascular pathology

PCSK9 overexpression combined with HFD feeding resulted in severe serum hypercholesterolemia and progressive weight gain over the 20-week experimental period in adult mice (Supplementary Figure 1). Marked elevation in serum cholesterol was most pronounced in the PCSK9+HFD group; however, PCSK9 overexpression alone also produced a modest but significant increase relative to control, whereas HFD alone produced only a small change. Body weight increased over time in all groups, with greater and comparable gains in the HFD-containing groups (HFD and PCSK9+HFD) relative to control and PCSK9 groups by the study endpoint.

We examined the effects of PCSK9 overexpression and HFD on cerebrovascular architecture. Vascular rarefaction, a reduction in the density of functional or structural microvessels, represents an early and functionally significant feature of CSVD[24, 25]. We assessed functional cerebrovascular architecture using lectin perfusion labeling combined with 2D immunofluorescence and 3D light-sheet microscopy. Lectin perfusion labels patent (perfused) vessels, enabling visualization of functional vascular networks. Quantitative morphometric analysis measured number of capillaries, number of junctions, and surface area ratio across cortex, striatum, and hippocampus (Figure 1A-D). Quantitative morphometric analysis demonstrated severe capillary rarefaction in PCSK9+HFD mice, with significantly reduced capillary density in cortex, striatum, and hippocampus compared to all other groups. HFD mice showed moderate reductions in vascular density (Fig 1A-D), while PCSK9 alone produced minimal effects, confirming synergistic effects of diet and PCSK9 overexpression. Three-dimensional reconstruction using light sheet microscopy demonstrated a sparser and less complex vascular network throughout the brain in PCSK9+HFD compared to controls (Figure 1E).

**Figure 1.**
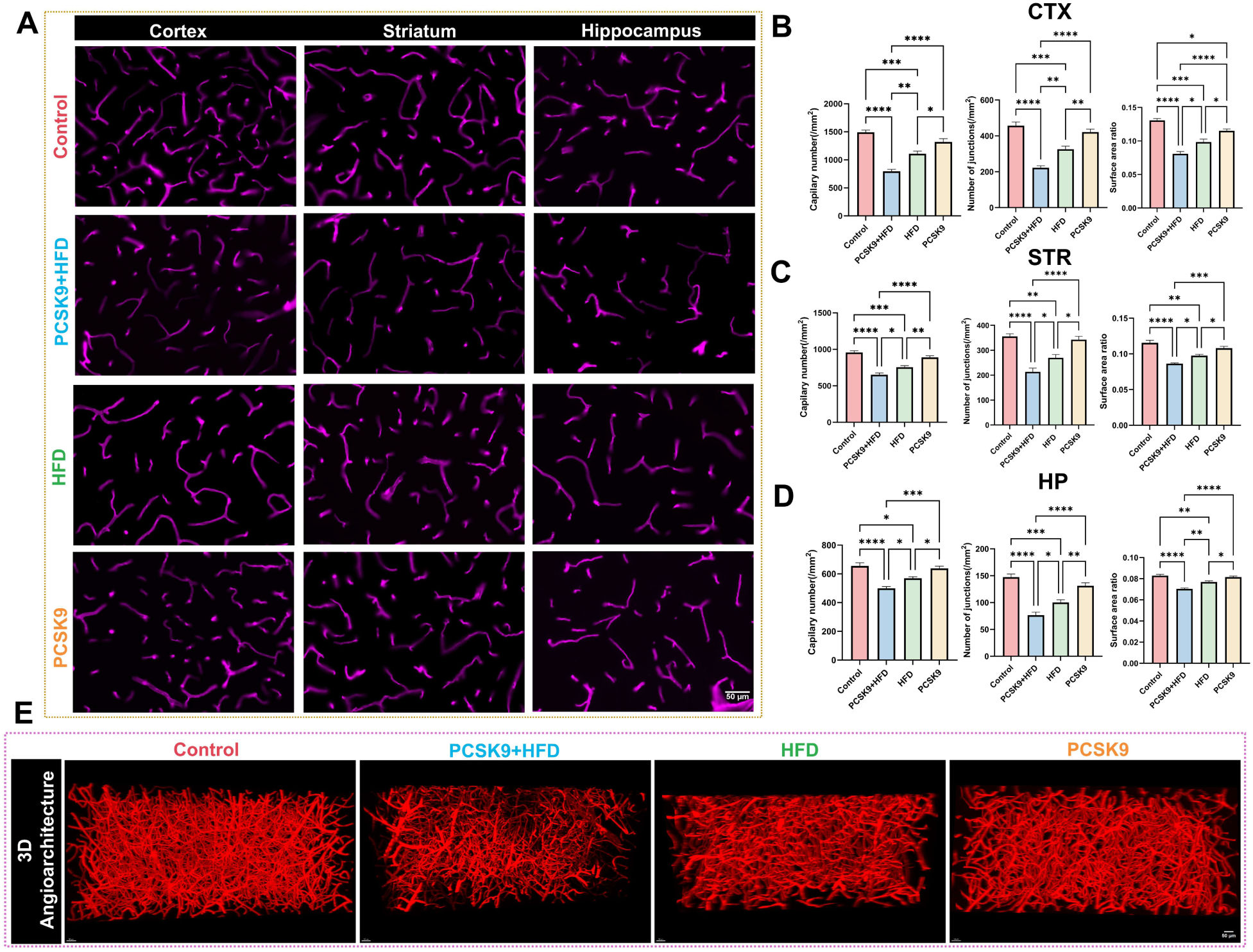
PCSK9 overexpression combined with high-fat diet induces profound cerebrovascular rarefaction and architectural disruption. (A) Representative confocal images of lectin-perfused (DyLight 649, magenta) brain sections showing microvascular architecture in cortex (CTX), striatum (STR), and hippocampus across treatment groups. Scale bars: 50 μm. (B) Quantitative morphometric analysis of vascular parameters in cortex showing capillary density (number/mm²), number of vascular junctions, and surface area ratio. (C) Corresponding analyses in striatum for capillary density, junction numbers, and surface area ratio. (D) Hippocampal vascular quantification including capillary density, junction numbers, and surface area ratio. (E) Three-dimensional light-sheet reconstructions of SHIELD/SmartClear-cleared whole brains: cortical renderings showing comprehensive vascular network disruption. Images show maximum intensity projections from control, PCSK9+HFD, HFD, and PCSK9 groups demonstrating progressive vascular rarefaction with combined treatment producing the most severe phenotype. Data represent mean ± SEM (n=4 per group). Statistical analysis by one-way ANOVA with Tukey’s post-hoc test. *P<0.05, **P<0.01, ***P<0.001, ****P<0.0001.

### PCSK9+HFD impairs neurovascular coupling and promotes amyloid-β accumulation

Neurovascular coupling, the relationship between neuronal activity and cerebral blood flow, is essential for proper brain function and is impaired in CSVD. Whisker stimulation is a well-established inducer of neuronal activation in the somatosensory cortex, providing a robust model for assessing neurovascular coupling[26]. Using laser speckle contrast imaging, we assessed cerebral blood flow responses to 30 seconds of whisker stimulation. Control mice exhibited a robust increase in cerebral blood flow during whisker stimulation. In contrast, PCSK9+HFD mice showed significantly attenuated flow responses, while HFD and PCSK9 mice displayed intermediate impairment (Figure 2).

**Figure 2.**
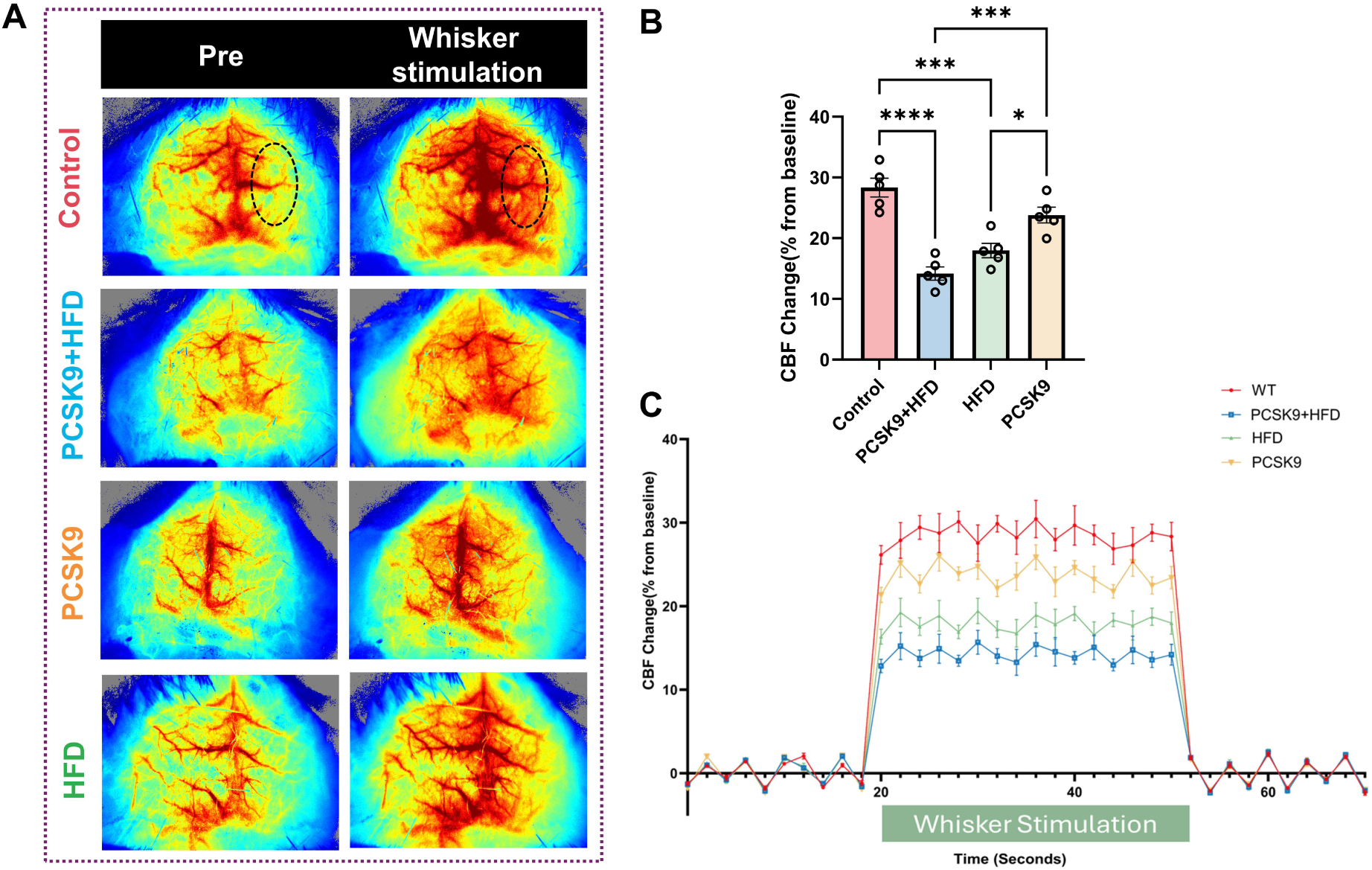
Impaired neurovascular coupling in PCSK9+HFD mice. (A) Representative laser speckle contrast images showing cerebral blood flow (CBF) patterns at baseline (Pre) and during whisker stimulation in anesthetized mice. Black oval demarcates the region of interest over the barrel cortex contralateral to whisker stimulation. Control mice show robust increase in cortical perfusion during stimulation, while PCSK9+HFD mice exhibit severely attenuated responses. (B) Quantification of CBF change (% from baseline) during 30-second whisker stimulation. PCSK9+HFD mice demonstrate significantly impaired neurovascular coupling compared to the control group. HFD mice show intermediate deficits, PCSK9 mice are comparable to controls. (C) Time course of CBF dynamics showing temporal profile of vascular responses. Control mice (red) display rapid CBF increase reaching ∼30% above baseline, while PCSK9+HFD mice (blue) show blunted response. Data shown as mean ± SEM (n=5 per group). One-way ANOVA with Tukey’s post-hoc test. **P<0.01, ****P<0.0001.

In addition to reduced capillaries (Fig 1) and blood flow (Fig 2), PCSK9+HFD mice developed extensive vascular pathology characteristic of CSVD. We found Aβ deposition within the walls of lectin-positive cortical microvessels, accompanied by scattered parenchymal deposits and perivascular clusters of CD68⁺ microglia/macrophages (Figure 3A). This vascular-wall localization of Aβ together with perivascular inflammatory cuffs resembles deposits found in human cerebral amyloid angiopathy (CAA), a well-recognized pathological feature of CSVD[27]. A similar pathology has been seen in genetic over-expression models of Alzheimer’s disease, human APOE4-5XFAD mouse model of Alzheimer’s disease [28]. To quantify vascular association and inflammatory response, we measured the area that was triple-positive for lectin, Aβ, and CD68 (Lectin⁺/Aβ⁺/CD68⁺) and double-positive for lectin and Aβ (Lectin⁺/Aβ⁺). Both metrics were significantly elevated in PCSK9+HFD mice compared with controls, with HFD alone showing intermediate increases and PCSK9 alone showing low levels (Figure 3B-C). The increase in Lectin⁺/Aβ⁺/CD68⁺ area indicates microglial recruitment around vascular Aβ.

**Figure 3.**
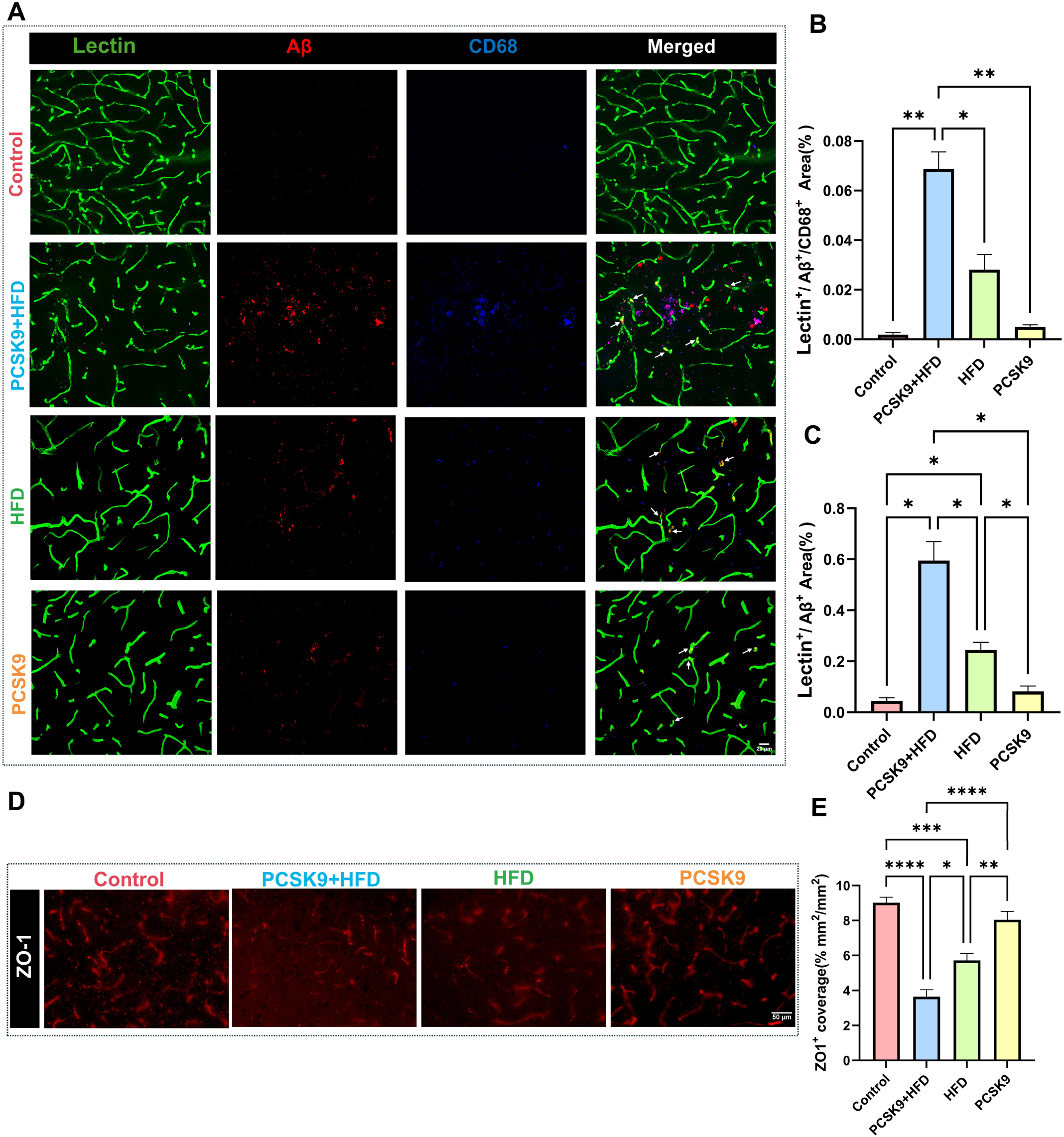
PCSK9 overexpression and HFD promote vascular pathology and blood-brain barrier disruption. **(A)** Triple immunofluorescence staining for lectin (green), amyloid-β (Aβ, red), and CD68 (blue) in cortical vessels. Control mice show patent vessels without amyloid deposition, while PCSK9+HFD mice exhibit extensive perivascular amyloid accumulation with CD68+ microglial activation surrounding affected vessels. HFD and PCSK9 groups show intermediate and mild pathology. Red arrows indicate triple-positive areas (lectin+/Aβ+/CD68+) representing amyloid-laden vessels with associated microglial activation. White arrows indicate dual-positive regions (lectin+/Aβ+) showing amyloid deposition. Scale bars: 20 μm. **(B)** Quantification of triple-positive area (lectin+/Aβ+/CD68+, %) demonstrating a pronounced convergence of vascular pathology, amyloid deposition, and neuroinflammation specifically in PCSK9+HFD mice. **(C)** Quantification of lectin+/Aβ+ area (%) showing amyloid deposition around vessels. **(D)** Representative images of ZO-1 immunostaining revealing tight junction integrity. Control mice display continuous ZO-1 expression along vessel walls, while PCSK9+HFD mice exhibit severely fragmented and reduced staining indicative of BBB breakdown. Scale bars: 50 μm. **(E)** Quantitative analysis of ZO-1 coverage confirming significant BBB disruption in PCSK9+HFD mice. Data represent mean ± SEM (n=4 per group). Statistical significance determined by Brown–Forsythe and Welch ANOVA, followed by Dunnett’s T3 post-hoc test for (B, C), and one-way ANOVA with Tukey’s post-hoc test for (E). *P<0.05, **P<0.01, ***P<0.001, ****P<0.0001.

BBB dysfunction is a key feature of CSVD. We assessed BBB integrity by examining the expression and coverage of the tight junction protein ZO-1 along cerebral vessels (Figure 3D-E). PCSK9+HFD mice exhibited significant disruption of ZO-1 coverage compared to controls. In control mice, ZO-1 showed continuous expression along vessel walls, whereas PCSK9+HFD mice displayed fragmented and reduced ZO-1 immunoreactivity, indicating compromised tight junction integrity. HFD mice showed intermediate reductions in ZO-1 coverage, while PCSK9 overexpression alone produced minimal effects on BBB integrity.

### PCSK9+HFD synergistically drive region**-**selective neuronal loss

Having established dysfunction and pathology, we next examined the consequences on neuronal atrophy. NeuN immunofluorescence revealed pronounced, region-dependent neuronal loss in PCSK9+HFD mice that significantly exceeded the reductions produced by either PCSK9 overexpression or HFD alone (Figure 4). In cortex (CTX), densely packed NeuN-positive nuclei observed in controls gave way to conspicuous cellular “voids,” indicating neuronal attrition. A similar pattern emerged along the hippocampal CA1 pyramidal layer, which is known to be sensitive to metabolic hypoperfusion[29], where neuronal rows appeared thinned and discontinuous. The striatum (STR) displayed an intermediate phenotype, consistent with its relatively greater resilience to chronic hypoperfusion reported in prior VCID models. Statistical comparison confirmed significant main effects of genotype, diet, and their interaction, underscoring a synergistic relationship between PCSK9 overexpression and high-fat feeding. Although sporadic CSVD classically involves deep gray nuclei and periventricular white matter, cortical microinfarcts and hippocampal atrophy are increasingly recognized as CSVD lesions linked to worse cognition[30-32].

**Figure 4.**
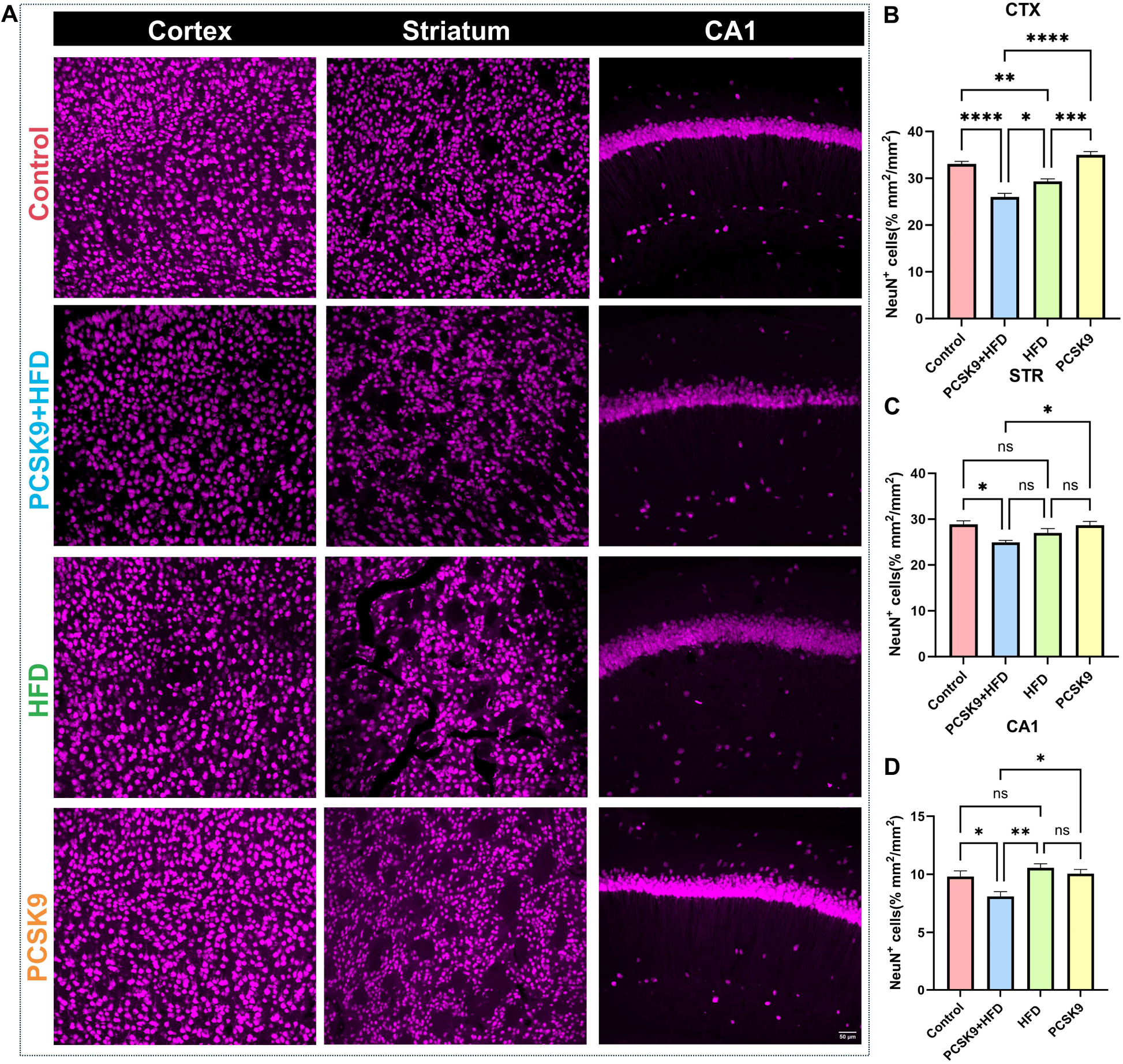
Regional patterns of neuronal loss in PCSK9+HFD mice. Representative NeuN immunofluorescence in cortex (CTX), striatum (STR), and hippocampal CA1 region across treatment groups. Control mice show densely packed neuronal nuclei in all regions. PCSK9+HFD mice exhibit significant neuronal loss characterized by reduced NeuN+ cell density and appearance of cellular "voids" particularly prominent in cortex and CA1. Striatum displays intermediate vulnerability. HFD and PCSK9 alone produce moderate and minimal neuronal loss. Scale bars: 50 μm. (B) Quantification of NeuN+ neuronal density in cortex showing significant reduction in PCSK9+HFD mice compared to all other groups. (C) Striatal neuronal counts revealing moderate but significant loss only in PCSK9+HFD group. (D) CA1 pyramidal neuron quantification demonstrating depletion in PCSK9+HFD mice. The vulnerability reflects differential metabolic demands and vascular supply across brain regions. Data represent mean ± SEM (n=4 per group). Statistical significance determined by one-way ANOVA with Tukey’s post-hoc test. *P<0.05, **P<0.01, ***P<0.001, ****P<0.0001; ns, not significant.

### PCSK9+HFD induces white matter injury and astrocyte reactivity

White matter injury is a prominent feature of CSVD pathology. To assess white matter integrity, we performed Luxol Fast Blue (LFB) staining across different brain regions (Figure 5A-D). PCSK9+HFD mice exhibited significant white matter damage characterized by reduced LFB staining intensity, particularly in the corpus callosum and external capsule, indicating myelin loss. Quantification of LFB staining revealed a substantial reduction in myelin content in PCSK9+HFD mice compared to controls. HFD mice showed moderate myelin loss in these regions, while PCSK9 mice displayed mild, localized reductions in LFB staining intensity.

**Figure 5.**
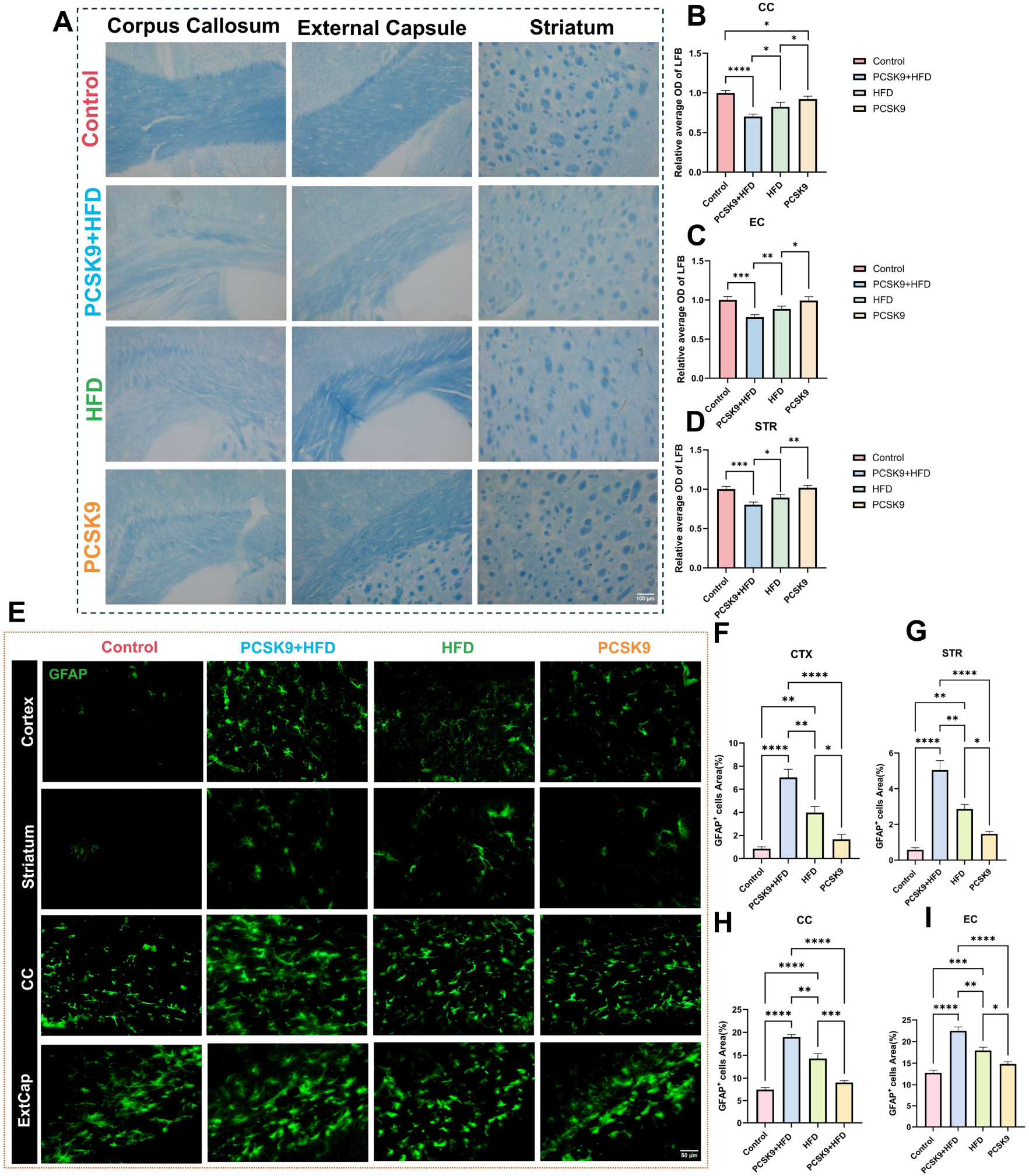
White matter injury and reactive astrogliosis in metabolic CSVD model. (A) Luxol Fast Blue (LFB) staining revealing myelin integrity in corpus callosum (CC), external capsule (EC), and striatum (STR). Control mice show intense blue staining indicating intact myelin. PCSK9+HFD mice display marked pallor and reduced staining intensity, particularly in CC and EC, indicating severe demyelination. HFD mice show moderate myelin loss while PCSK9 alone produces minimal changes. Scale bars: 100 μm. (B-D) Quantification of relative LFB optical density (normalized to control) in CC (B), EC (C), and STR (D), confirming significant white matter injury in PCSK9+HFD mice. (E) GFAP immunofluorescence (green) demonstrating astrocyte reactivity across brain regions. PCSK9+HFD mice exhibit extensive astrogliosis with hypertrophic astrocytes throughout white and gray matter. Scale bars: 50 μm. (F-I) Quantitative analysis of GFAP+ area (%) in cortex (F), striatum (G), corpus callosum (H), and external capsule (I). Data shown as mean ± SEM (n=4 per group). One-way ANOVA with Tukey’s post-hoc test. *P<0.05, **P<0.01, ***P<0.001, ****P<0.0001.

Astrocytes play essential roles in maintaining neurovascular unit integrity and respond to various pathological stimuli. GFAP immunostaining revealed substantial astrocyte reactivity in PCSK9+HFD mice across all brain regions examined (Figure 5E-I). The GFAP-positive area was significantly increased in the corpus callosum, external capsule, cortex, and striatum of PCSK9+HFD mice compared to controls.

### PCSK9+HFD promotes microglial activation and lipid droplet accumulation

Neuroinflammation orchestrated by activated microglia is a central driver of CSVD pathogenesis[33, 34]. To comprehensively characterize microglial responses to metabolic stress, we combined high-resolution confocal microscopy with light sheet microscopy. Iba1 immunofluorescence revealed profound microglial activation in PCSK9+HFD mice, with dramatically increased cell density. HFD mice showed a moderate increase in microglial density, while PCSK9 mice exhibited a modest increase (Figure 6A-B). Beyond proliferation, microglia in PCSK9+HFD mice exhibited striking morphological transformation. Three-dimensional reconstructions of individual cells revealed a transition from the ramified, homeostatic morphology observed in controls to activated phenotypes characterized by the presence of amoeboid-like cells and hyper-ramified cells with increased process complexity[35]. These hypertrophic amoeboid and hyper-ramified Iba1+ cells were often concentrated near altered vasculature.

**Figure 6.**
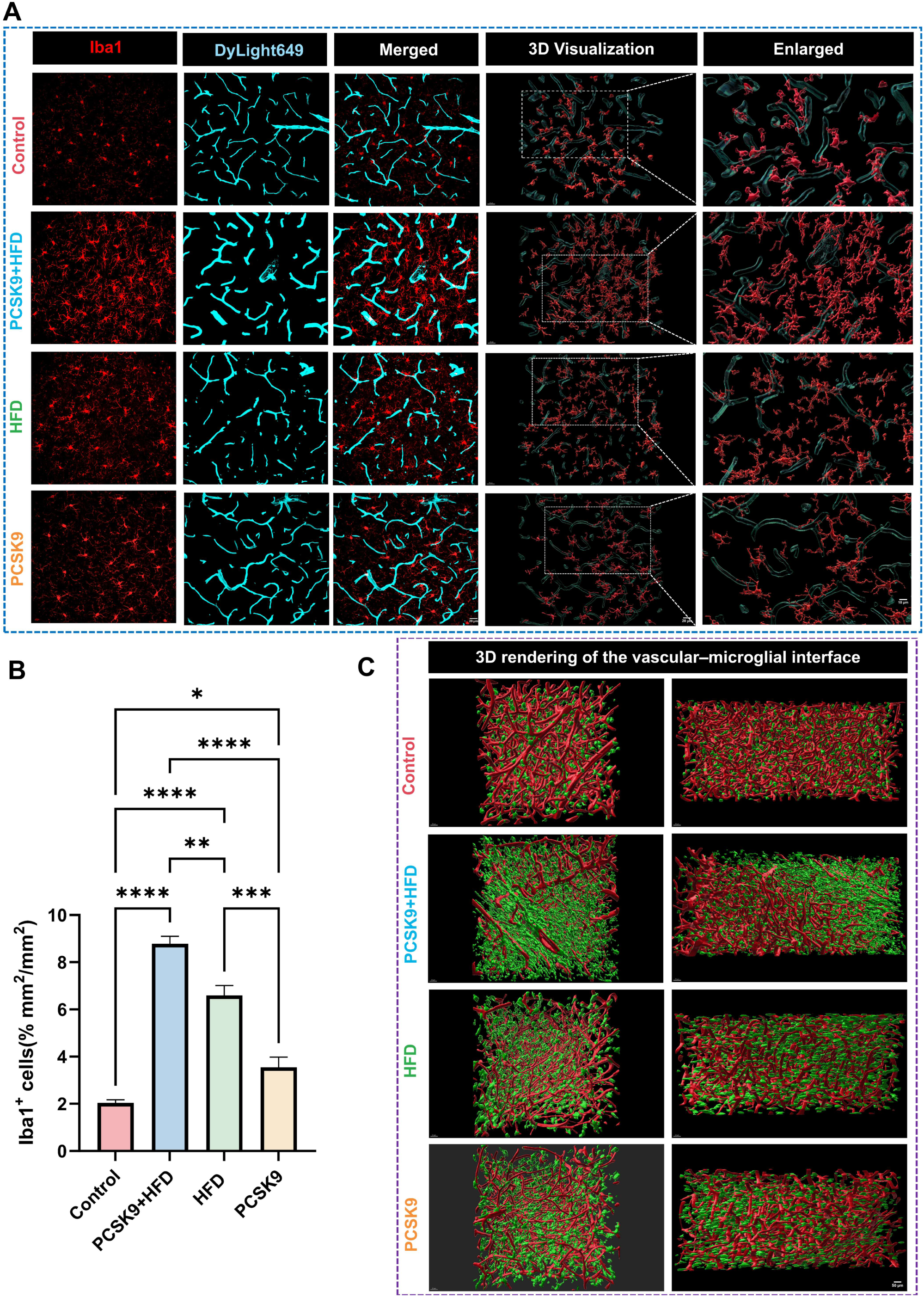
Microglial activation and perivascular clustering in PCSK9+HFD mice. (A) Representative confocal images showing spatial relationship between Iba1+ microglia and lectin-labeled vessels. Control mice display microglia with uniform distribution. PCSK9+HFD mice show dramatic microglial activation with amoeboid morphology and perivascular clustering. Three-dimensional visualization reveals microglia forming aggregates around vessels. Enlarged panels demonstrate individual microglial morphology and vascular contacts. Scale bars: Iba1/DyLight649/Merged and 3D Visualization, 20 µm; Enlarged, 10 µm. (B) Quantification of Iba1+ cell density showing significant microglial proliferation in PCSK9+HFD mice. (C) Three-dimensional rendering of vascular-microglial interface from light sheet microscopy. Control mice maintain sparse microglial distribution with limited vascular contact (red: vessels, green: microglia). PCSK9+HFD mice exhibit extensive microglial infiltration and perivascular accumulation, with microglia extending multiple processes to ensheath blood vessels. Scale bars: 50 μm. Data represent mean ± SEM (n=4 per group). One-way ANOVA with Tukey’s post-hoc test. *P<0.05, **P<0.01, ***P<0.001, ****P<0.0001.

The spatial relationship between activated microglia and compromised vasculature revealed a striking pattern of cellular reorganization. Light sheet microscopy with three-dimensional reconstruction uncovered that microglia in PCSK9+HFD mice formed dense perivascular clusters (Figure 6C). In control mice, microglia maintained their characteristic uniform distribution with vascular contact, whereas PCSK9+HFD mice showed microglia extending multiple processes to ensheath blood vessels. This close association suggests active surveillance or responses to vascular damage. CD68 immunostaining, a marker of microglial phagocytic activity, was dramatically increased in PCSK9+HFD mice, indicating enhanced microglial activation (Figure 7A-B). CD68 expression was moderately increased in HFD mice and minimally elevated in PCSK9 mice compared to controls. Sustained phagocytic activation may contribute to a chronic neuroinflammatory state.

**Figure 7.**
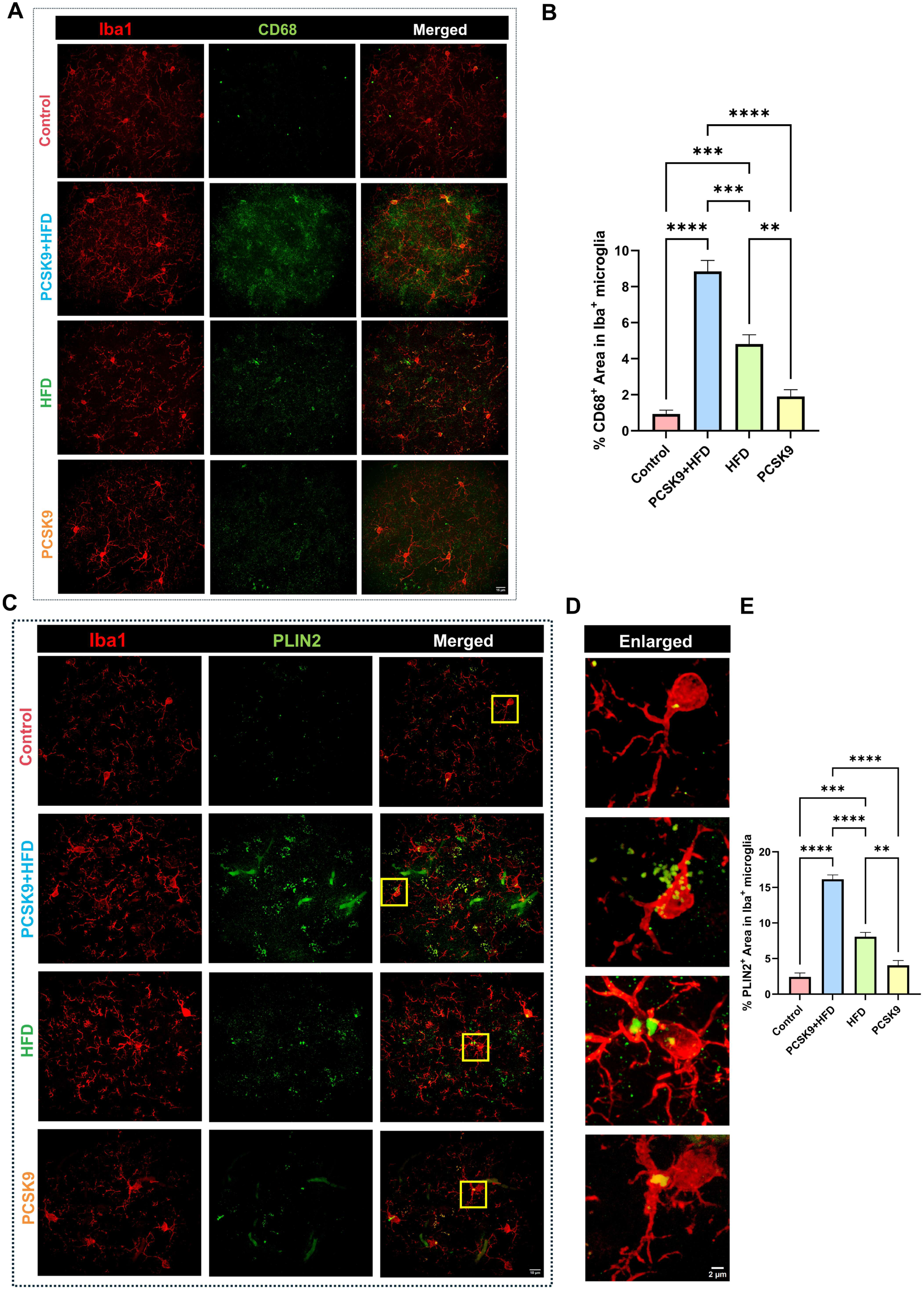
Microglial activation and emergence of lipid-droplet accumulating microglia (LDAM) in PCSK9+HFD mice. **(A)** Co-immunostaining for Iba1 (red) and CD68 (green) revealing microglial phagocytic activity. Control mice show minimal CD68 expression. PCSK9+HFD mice display extensive CD68+ staining within Iba1+ microglia, indicating sustained phagocytic activation. HFD mice show moderate activation while PCSK9 alone produces minimal changes. Scale bars: 15 μm. **(B)** Quantification of CD68+ area within Iba1+ microglia (%) demonstrating significantly enhanced phagocytic activity in PCSK9+HFD mice. **(C)** Co-immunostaining for Iba1 (red) and perilipin-2 (PLIN2, green), a specific marker of lipid droplet surface proteins. Control mice show minimal PLIN2 expression in microglia. PCSK9+HFD mice exhibit dramatic PLIN2 accumulation within Iba1+ cells, indicating extensive lipid droplet formation. HFD alone induces moderate LDAM formation, while PCSK9 produces minimal effect. Scale bars: 10 μm. **(D)** High-magnification images from panel C showing detailed morphology of PLIN2+ lipid droplets within individual Iba1+ microglia. Control microglia show minimal lipid content. PCSK9+HFD microglia are laden with abundant intracellular PLIN2+ lipid droplets. Scale bars: 2 μm. **(E)** Quantification of PLIN2+ area in Iba1+ microglia (%) confirming synergistic effect of PCSK9 and HFD on microglial lipid accumulation. Data represent mean ± SEM (n=4 per group). One-way ANOVA with Tukey’s post-hoc test. **P<0.01, ***P<0.001, ****P<0.0001.

Strikingly, we observed the emergence of a distinctive lipid-droplet accumulating microglial (LDAM) phenotype. To characterize microglial lipid metabolism in this model, we performed co-immunostaining for perilipin-2 (PLIN2), a specific marker of lipid droplet surface proteins (Figure 7C,E). PLIN2 expression was dramatically elevated in microglia of PCSK9+HFD mice. HFD alone induced moderate LDAM formation, while PCSK9 monotherapy produced minimal effect, suggesting a synergistic effect of hypercholesterolemia and high-fat diet on microglial lipid uptake. High-magnification images revealed abundant intracellular lipid accumulation in microglia of PCSK9+HFD mice (Figure 7D). These LDAM exhibited distinctive activated morphologies, particularly in areas proximal to altered vasculature. This morphological signature, combined with their localization near compromised vessels, suggests that the increase in LDAMs reflects the role of perivascular microglia in responding to hypercholesterolemia. Excessive lipid loading is known to disrupt lysosomal flux and limit microglial phagocytic competence, providing a mechanistic rationale for the persistence of perivascular amyloid despite elevated CD68 expression. Taken together, these findings suggest that the combination of hypercholesterolemia and HFD synergistically drive LDAM formation, representing a context-specific metabolic stress response that links systemic metabolic dysfunction to cerebrovascular injury in CSVD.

### PCSK9+HFD impairs cognitive function and alters behavior

To assess the functional consequences of cerebrovascular and neuroinflammatory pathology, we performed open field and novel object recognition (NOR) testing. In the open field test, PCSK9+HFD mice spent significantly less time in the center zone compared to controls, indicating increased anxiety-like behavior. HFD mice also showed reduced center time at a level comparable to PCSK9+HFD mice, while PCSK9 mice displayed a less pronounced reduction (Figure 8A-E). Total distance traveled and average movement speed did not differ significantly between groups. The NOR test was used to evaluate recognition memory. In the NOR test, PCSK9+HFD mice exhibited severe recognition memory impairment (Figure 8F-H). The discrimination index was significantly reduced in PCSK9+HFD mice compared to controls. In particular, PCSK9+HFD mice spent proportionally more time with the familiar object, indicating marked recognition-memory impairment. These functional assessments confirm that PCSK9+HFD-induced cerebrovascular pathology translates to meaningful cognitive and behavioral impairments characteristic of CSVD.

**Figure 8.**
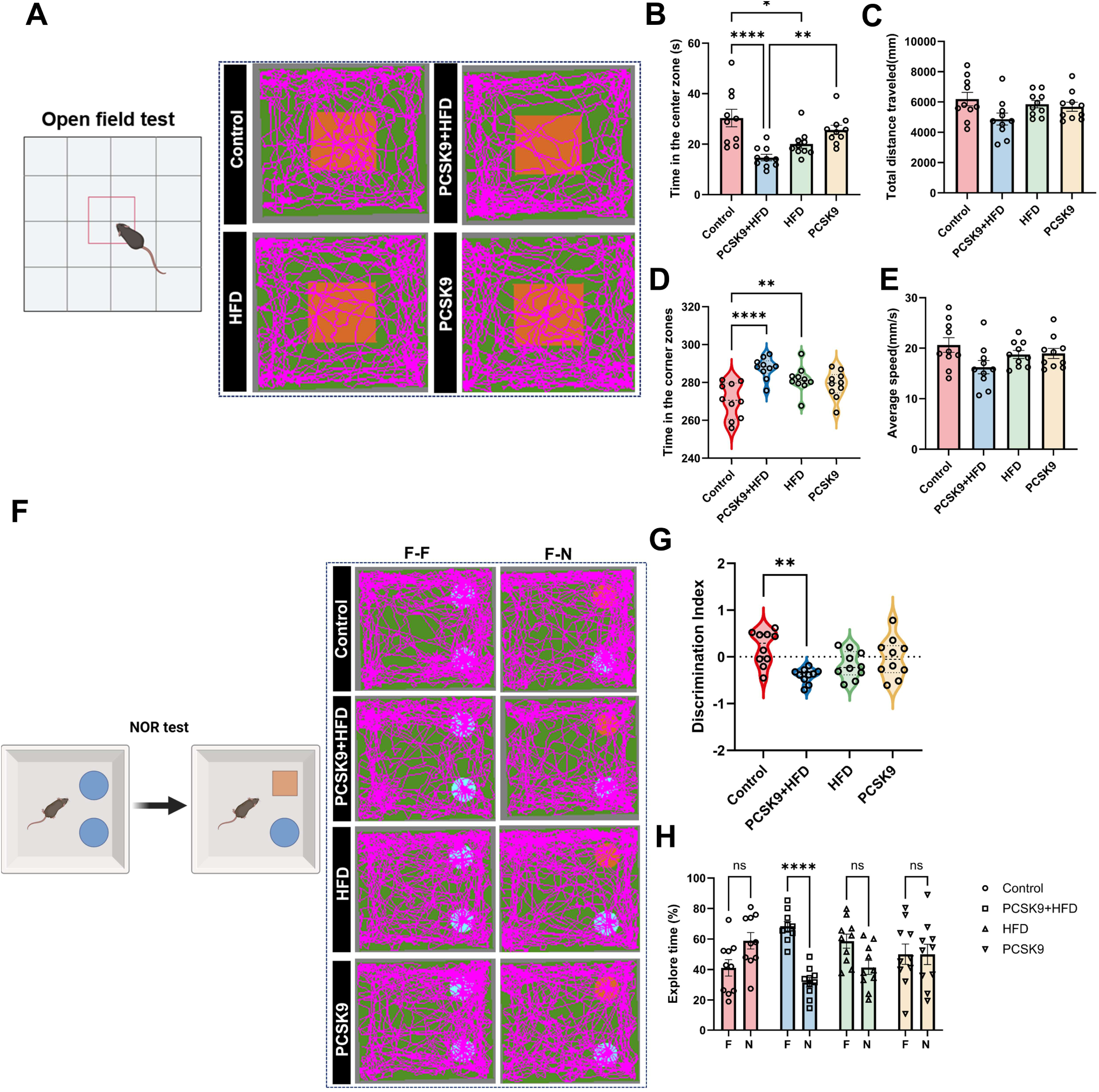
Cognitive impairment and behavioral alterations following cerebrovascular pathology. **(A)** Representative movement traces from OFT showing exploration patterns. Control mice explore the entire arena including center zone (red square). PCSK9+HFD and HFD mice display thigmotactic behavior, avoiding center and preferring periphery. (B) Time in center zone quantification revealing increased anxiety-like behavior in PCSK9+HFD and HFD groups. (C) Total distance traveled showing preserved locomotor function across all groups. (D) Time spent in corner zones (violin plot) further supporting anxiety phenotype, with PCSK9+HFD mice spending significantly more time in corners. (E) Average movement speed confirming absence of motor deficits. (F) Representative exploration patterns from NOR test showing interaction with familiar (F) and novel (N) objects. Control mice show preferential exploration of the novel object, while PCSK9+HFD mice show impaired novelty discrimination with reduced preference for the novel object. (G) Discrimination index calculated as (time with novel – time with familiar)/(total exploration time). PCSK9+HFD mice show significantly reduced discrimination index indicating severe recognition memory impairment. (H) Exploration time (%) for each object reveals that control mice spend more time exploring the novel object compared to the familiar object (demonstrating intact recognition memory), while PCSK9+HFD mice spend significantly more time with familiar object and less with novel object, demonstrating reversal of normal exploration pattern. Data shown as mean ± SEM (n=10 per group). Statistical analysis by one-way ANOVA with Tukey’s post-hoc test for (B-E, G); two-way ANOVA with Šidák’s post-hoc test for (H). *P<0.05, **P<0.01, ****P<0.0001; ns, not significant.

## Discussion

This study establishes a comprehensive mouse model that recapitulates a spectrum of human CSVD pathology through the synergistic effects of hepatic PCSK9 overexpression and HFD. Our findings demonstrate that metabolic dysfunction drives a cascade of cerebrovascular alterations encompassing vascular rarefaction, neurovascular uncoupling, BBB disruption, white matter injury, neuroinflammation, neuronal loss, and cognitive impairment. The convergence of these pathological features within a 20-week timeframe offers a valuable platform for understanding CSVD pathogenesis.

A critical finding of our study is that severe cerebrovascular pathology requires synergistic interactions between PCSK9-mediated hypercholesterolemia and HFD. PCSK9 overexpression alone elevates serum cholesterol modestly but remains below the threshold necessary to induce significant cerebrovascular pathology. This threshold-dependent mechanism has translational relevance as individuals with genetic variants impairing cholesterol metabolism (PCSK9 gain-of-function, LDLR deficiency) may remain relatively asymptomatic on low-cholesterol diets but develop accelerated cerebrovascular disease with high dietary intake.

The profound reduction in cerebrovascular density observed in PCSK9+HFD mice represents a fundamental alteration in brain vascular architecture[36]. Recent studies have established vascular rarefaction as an early and critical feature of CSVD that precedes other pathological changes[37]. The mechanisms driving vascular loss likely involve multiple pathways: direct endothelial toxicity from hypercholesterolemia[38], reduced VEGF signaling, and impaired angiogenesis[39]. Importantly, the synergistic effect of PCSK9 and HFD on vascular density suggests that multiple metabolic hits are required to overcome compensatory mechanisms maintaining vascular homeostasis[40]. Prior studies in CSVD models have implicated chronic hypoperfusion as triggering a cascade of cellular responses including HIF-1α activation, metabolic stress, and ultimately cellular dysfunction[41]. Future studies in this model will further examine the endothelial cellular events accompanying vascular disruption as well as delineate the timing of vascular loss and other cerebral pathologies.

The severe impairment of neurovascular coupling in PCSK9+HFD mice represents a critical functional deficit that may directly contribute to white matter injury and neuronal loss and underlie cognitive deficits. Neurovascular coupling ensures that neuronal activity is matched by appropriate increases in local blood flow, a process essential for meeting the high metabolic demands of active brain regions[42]. Multiple mechanisms likely contribute to neurovascular uncoupling in our model. Endothelial dysfunction, evidenced by BBB disruption and altered tight junction expression, impairs NO-mediated vasodilation[43]. Pericyte dysfunction, suggested by altered smooth muscle actin patterns, disrupts capillary flow regulation[44]. Astrocyte end-feet swelling and reactive gliosis interfere with gliovascular signaling[45]. Together, these changes disrupt normal neurovascular unit function in response to neural activity, leading to activity-dependent hypoxia and metabolic stress. Clinically, neurovascular uncoupling is increasingly recognized as an early biomarker of CSVD that precedes structural changes on conventional MRI[46]. Recapitulation of this functional deficit reinforces relevance of the PCSK9+HFD model for understanding human disease.

Extensive BBB disruption as evidenced by fragmented ZO-1 expression in PCSK9+HFD mice represents a pivotal event in CSVD pathogenesis. BBB disruption in our model also likely results from convergent insults. Hypercholesterolemia directly damages endothelial cells through oxidative stress and inflammatory signaling[47], while HFD-induced systemic inflammation amplifies endothelial dysfunction through circulating inflammatory mediators[48]. BBB disruption allows for extravasation of blood-borne proteins including fibrinogen and albumin triggering perivascular inflammation and edema[49]. Extravasated proteins form deposits that impair interstitial fluid drainage, contributing to waste accumulation[50]. The disrupted BBB also permits entry of peripheral immune cells and circulating lipids, fueling neuroinflammation and LDAM formation[51].

A particularly striking finding is the emergence of vascular Aβ deposition in wild-type mice without genetic Alzheimer’s disease mutations. Classical CAA models typically require APP overexpression or familial AD mutations to drive vascular amyloid accumulation[52]. In contrast, PCSK9+HFD mice develop CAA-like pathology purely under metabolic stress on a non-transgenic background. Remarkably, LDAM and vascular Aβ pathology appear in a cerebral milieu of hypercholesterolemia with BBB vulnerability. While our data do not establish direct causality, prior work shows that microglial lipid accumulation impairs Aβ handling and that reducing microglial lipid load can improve Aβ clearance in transgenic models[53]. These findings raise the possibility that metabolic stress-induced microglial lipid loading may contribute to impaired Aβ clearance, creating conditions permissive for vascular amyloid accumulation.

Additional mechanisms likely include access of peripheral Aβ sources when barriers are compromised[54], as well as local Aβ production by stressed vascular cells[55]. The emergence of CAA-like pathology in wild-type mice therefore suggests that metabolic syndrome contributes to cerebrovascular amyloid pathogenesis and motivates testing aggressive lipid-lowering or microglial lipid–lysosomal interventions.

Our findings support the emergence of LDAM in PCSK9+HFD mice. Consistent with prior reports of LDAMs[56, 57], we observed an increased frequency of microglia with evidence of lipid inclusions. Prior work has linked these features to aging and neurodegeneration[57-61]. Importantly, the PCSK9+HFD model recapitulates these changes in the context of diet-induced metabolic syndrome and hypercholesterolemia, supporting the idea that systemic dyslipidemia may accelerate LDAM formation and contribute to microglial dysfunction in aging and VCID. Given that LDAM are increasingly recognized in human aging and Alzheimer’s disease, our findings highlight PCSK9+HFD mice as a model in which microglial lipid dysregulation and cerebrovascular pathology converge. The mechanistic basis for LDAM formation likely involves multiple converging pathways. First, BBB disruption in PCSK9+HFD mice permits entry of circulating lipids and lipoproteins into brain parenchyma, creating a lipid-rich microenvironment[62]. Second, impaired cholesterol efflux due to APOE dysfunction may trap lipids within microglia[63]. Third, overwhelmed lysosomal degradation capacity could lead to lipid droplet accumulation[64]. Recent evidence suggests that microglial lipid metabolism directly regulates inflammatory responses through metabolic reprogramming, positioning LDAM at the intersection of metabolism and neuroinflammation[65]. Targeting LDAM could thus represent a novel therapeutic strategy for metabolic CSVD.

CD68 immunostaining indicated increased phagocytic activity in PCSK9+HFD microglia, yet this activation appeared maladaptive, as perivascular amyloid persisted despite robust CD68 expression [53, 66]. Several mechanistic questions warrant future investigation. The precise signals driving LDAM formation in metabolic syndrome remain unclear and may include modified lipoproteins, damage-associated molecular patterns, or metabolic stress signals. Future studies examining functional heterogeneity within LDAM populations will also provide valuable insights into the contribution of this cell type to pathology. The contribution of peripheral immune cells versus resident microglia to the LDAM pool also needs clarification. Taken together, targeting LDAM could be a viable therapeutic strategy to combat neuroinflammatory consequences of metabolic syndrome and hypercholesterolemia.

Downstream of vascular and inflammatory changes, PCSK9+HFD mice exhibited both white matter injury and neuronal loss. White matter showed severe myelin loss in corpus callosum and external capsule, mirroring the characteristic pattern of human CSVD[67]. White matter’s high metabolic demands and limited collateral circulation render it particularly vulnerable to hypoperfusion[68]. Our observation that association fibers are more affected than projection fibers aligns with DTI studies in human CSVD showing preferential disruption of inter-hemispheric and association pathways[69]. The mechanisms of white matter injury in our model likely involve both ischemic and inflammatory components. Chronic hypoperfusion due to vascular rarefaction creates energy failure in oligodendrocytes, leading to myelin breakdown[70]. Inflammatory mediators from activated microglia and astrocytes directly damage myelin and prevent remyelination[71]. The accumulation of myelin debris further activates microglia, creating a self-perpetuating cycle of white matter damage[72]. Importantly, white matter injury serves as the anatomical substrate for cognitive impairment in CSVD[73]. Disruption of white matter tracts disconnects cortical-subcortical circuits essential for cognitive processing[74]. Our behavioral data showing impaired recognition memory and increased anxiety directly correlate with the anatomical distribution of white matter pathology, supporting the concept of CSVD as a disconnection syndrome[75].

Neuronal loss, which was especially prominent in metabolically active regions like the hippocampus, further reflected the interplay between metabolic demand and vascular supply. Multiple mechanisms likely contribute to neuronal death in our model. Chronic hypoperfusion leads to energy failure and excitotoxicity[76]. Loss of trophic support from dysfunctional astrocytes and microglia deprives neurons of essential survival factors[77, 78]. Accumulation of toxic metabolites due to impaired clearance creates a hostile microenvironment[79] [80]. Functionally, recognition memory deficits and anxiety-like behaviors in PCSK9+HFD mice correspond to these anatomical patterns. The selective impairment of recognition memory, mild with either PCSK9 overexpression or HFD alone, but severely affected by combined PCSK9+HFD, suggests a threshold of pathophysiological effects that ultimately converge to affect cognitive reserve[81]. This pattern aligns with clinical observations that cognitive symptoms emerge only after accumulation of significant pathological burden.

Comparison with existing CSVD models highlights unique advantages of our approach. Genetic (monogenic) models recapitulate specific human arteriopathies: NOTCH3/CADASIL[82] and knock-in or transgenic mice that develop granular osmiophilic material and mural cell pathology [83-85], COL4A1/2 mutations causing basement membrane fragility[86], and TgSwDI mice with vasculotropic amyloid[87]; and illuminate specific monogenic pathways, but represent a small minority of human stroke/CSVD presentations and lack metabolic components. Non-genetic approaches such as bilateral carotid artery stenosis (BCAS) produce robust white matter injury and cognitive deficits within weeks, but lacks the metabolic risk factors central to human CSVD [88]. Spontaneously hypertensive stroke-prone rats (SHRSP), a commercially available strain developed through selective inbreeding, spontaneously develop hypertensive arteriopathy characterized by arteriolar lipohyalinosis, blood-brain barrier breakdown, accelerated by salt-loading[89, 90]. However, SHRSP primarily model hypertensive arteriopathy rather than the full spectrum of metabolic CSVD, and are frequently used as a sensitized background for experimental stroke studies[91] rather than for spontaneous lacunar pathology. Diet-induced models require prolonged exposure for mild phenotypes[92]. Our PCSK9+HFD model uniquely combines human-relevant dyslipidemia with dietary metabolic stress in wild-type mice, converging on key CSVD features such as microvascular rarefaction, neurovascular uncoupling, BBB leakage, LDAM, and CAA-like vascular Aβ, thus bridging the gap between monogenic precision and sporadic disease complexity.

Several limitations warrant consideration. First, while our model recapitulates many features of human CSVD, the accelerated progression may not fully reflect all aspects of chronic human pathology. Second, we focused on male mice, but acknowledge sex differences in CSVD pathogenesis are likely to be significant and require future investigation in female mice. Third, while we demonstrate correlation between LDAM and cognitive impairment, establishing causal and mechanistic relationships will require further study including selective LDAM depletion or experiments targeting microglial lipid pathways, such as CSF1R inhibition[93] or TREM2 modulation[94].

In conclusion, we establish hepatic PCSK9 overexpression combined with HFD as a driver of accelerated CSVD-like pathology and identify LDAM as a novel cellular phenotype linking metabolic dysfunction to cerebrovascular disease. This model recapitulates key features of human CSVD including vascular rarefaction, BBB disruption, white matter injury, neuroinflammation, and cognitive impairment within an accelerated timeframe. The discovery of the preponderance of LDAMs in this model provides new insights into how metabolic stress transforms protective microglia into contributors of pathology. Our findings suggest that targeting PCSK9 and microglial lipid metabolism may offer therapeutic opportunities for preventing or treating CSVD. This work establishes a valuable platform for mechanistic studies and therapeutic development while highlighting the critical importance of addressing metabolic risk factors in preserving cerebrovascular health during aging.

## Funding acknowledgments

This work was supported by pilot funds from the Hellman Family fund, UCSF Nutrition Obesity Research Center (NORC; P30DK09872), and UCSF Research Evaluation and Allocation Committee. NSS is supported by 1IK2BX005369. ELG is supported by DP2AI175641 and by a Pilot & Feasibility Award from the UCSF NORC (P30 DK098722). Additional support was provided by the Bakar Aging Research Institute (awards to ELG and AL).

