## Supplementary Figure1 for "PCSK9 and High-Fat Diet Synergistically Induce Neurovascular Dysfunction and Neuroinflammation"

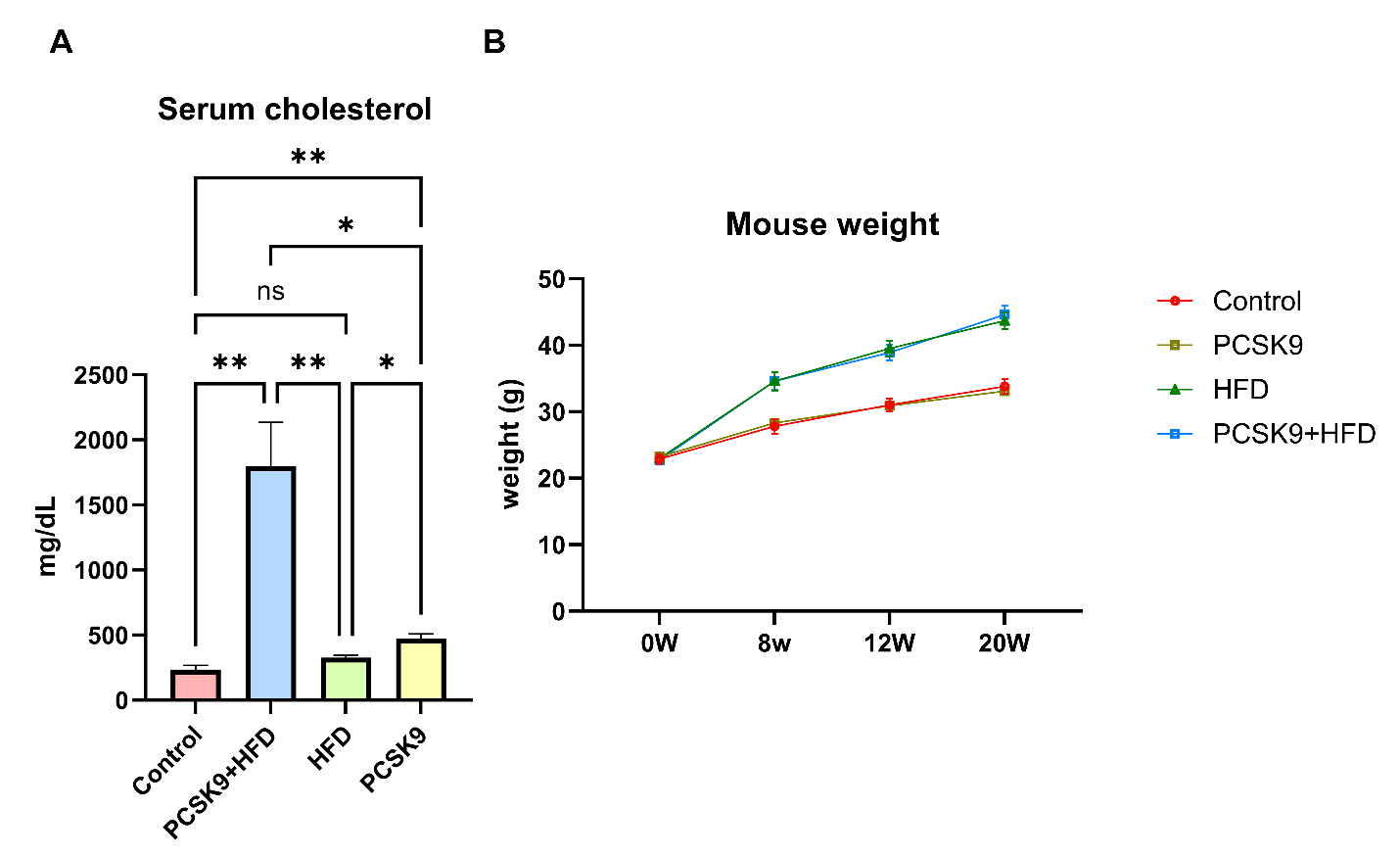

**Supplementary Figure 1. Systemic lipid and weight trajectory in PCSK9 overexpression and high-fat diet model.** (A) Serum total cholesterol at study endpoint (week 20). PCSK9+HFD shows the expected marked hypercholesterolemia relative to Control, PCSK9 alone, and HFD alone. (B) Body weight trajectory from baseline (0 w) to week 20. Data shown as mean ± SEM (n=10 per group). Statistical analysis by Brown–Forsythe and Welch ANOVA, followed by Dunnett's T3 post-hoc test. *P<0.05, **P<0.01, ns, not significant.
